# An open-source and low-cost feeding system for zebrafish facilities

**DOI:** 10.1101/558205

**Authors:** Astou Tangara, Gérard Paresys, Firas Bouallague, Yvon Cabirou, Jozsua Fodor, Victor Llobet, Germán Sumbre

## Abstract

Zebrafish is an established animal model used in the fields of developmental biology and genetics for more than 60 years, and among the first models to become genetically tractable. More recently, zebrafish has also become a model of reference for human diseases and systems neuroscience.

The current extensive use of zebrafish in research and pharmaceutical companies promoted the development of several commercial husbandry systems specially designed for zebrafish. However, feeding is still a challenging and arduous task that can result in occupational disorders of the personnel working at the zebrafish facilities (e.g. tendinitis, back pain, etc.). To palliate these risks, a commercial robotic approach has been developed, yet its expensive cost makes this solution accessible only to very large fish facilities. Most mid-size and small facilities with limited resources still use manual feeding methods.

Here, we propose two custom-made open-source semi-automatic feeding systems for dry and live food, capable of preventing and/or alleviating occupational disorders, improve feeding accuracy and decrease feeding time. Both systems are designed for mid-size or small fish facilities. They are cheap and can be easily and rapidly built using 3D printing and standard electronic and lab components.

## Introduction

Adequate feeding, monitoring and taking care of zebrafish is essential for their health and well-being. For an optimal nutrition it is important to use different types of food, both live (e.g. Artemia) and dry (*e.g. MG 300*), and regulate the distribution to avoid over or underfeeding.

In mid-size or small zebrafish facilities, fish are fed two to five times a day. This task is assured by dedicated personnel usually using manual approaches that demand repetitive motor gestures (e.g. forceps-like movement when delivering dry food or squeezing a wash bottle when feeding with live food). In addition, during feeding the arm remains in an extended position for long periods of time. At the long-term, these constraints can result in occupational disorders such as tendinopathy, affection of the rotator cuff at the shoulders, and tendinitis or tenosynovitis at the level of the hand [1, 2].

For optimizing fish feeding, and to solve and prevent occupational disorders of the personnel, we developed two semi-automatic feeding systems. One designed for live food and the other for dry food.

These systems were specifically conceived to reduce the feeding time, to improve food-delivery accuracy (delivering the right quantity of food), but most importantly to take into account ergonomic constraints : 1) decrease in the number repetitive gestures, 2) a better distribution of weight, 3) a better grip and 4) a better work posture.

These systems are easy to build, inexpensive and open-source so users can improve them or modify them to their own needs or constraints. In this article, we provide all the required information for their replication.

## Results

### Live-food dispenser system

In our fish facility, we feed zebrafish once a day using larvae of Artemia salina as live food. Artemia were delivered into the aquariums using a washing bottle which after long-term use can cause an inflammation in the flexor tendons of the fingers. To prevent and remedy this problem, we designed an effortless semi-automatic system for the accurate distribution of live food (Figure 1).

**Figure 1:**
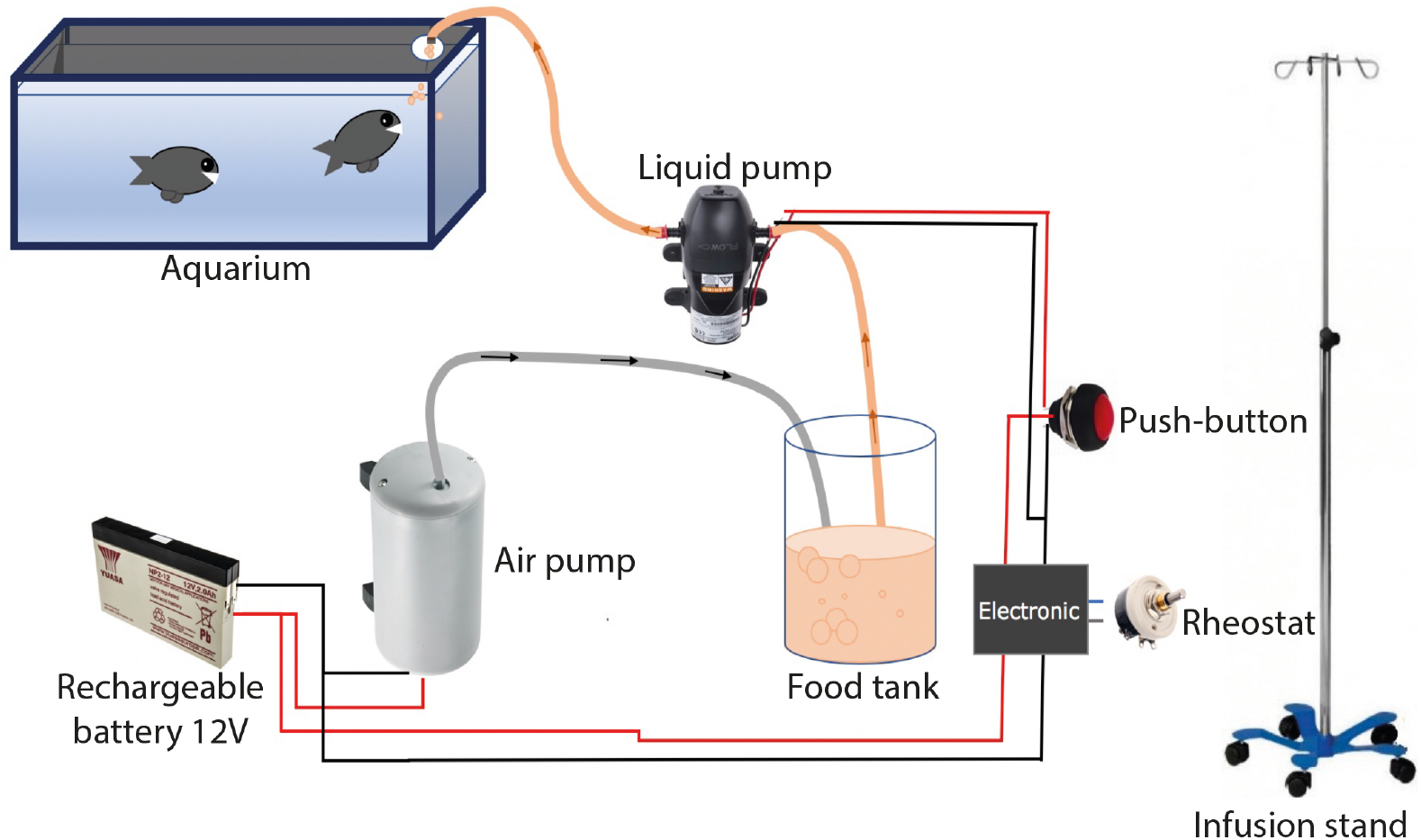
Scheme of the live-food dispenser system.

This system is based on an electric pump (*Xylem Flojet, RLFP122202D*) capable of sucking live Artemia and deliver them to the fish tanks (Figure 2). The rate of delivered Artemia can be adjusted by regulating the pump’s speed (Figure 3). The pump is trigger by a sensitive and light-weight button held by the person in charge of feeding.

**Figure 2:**
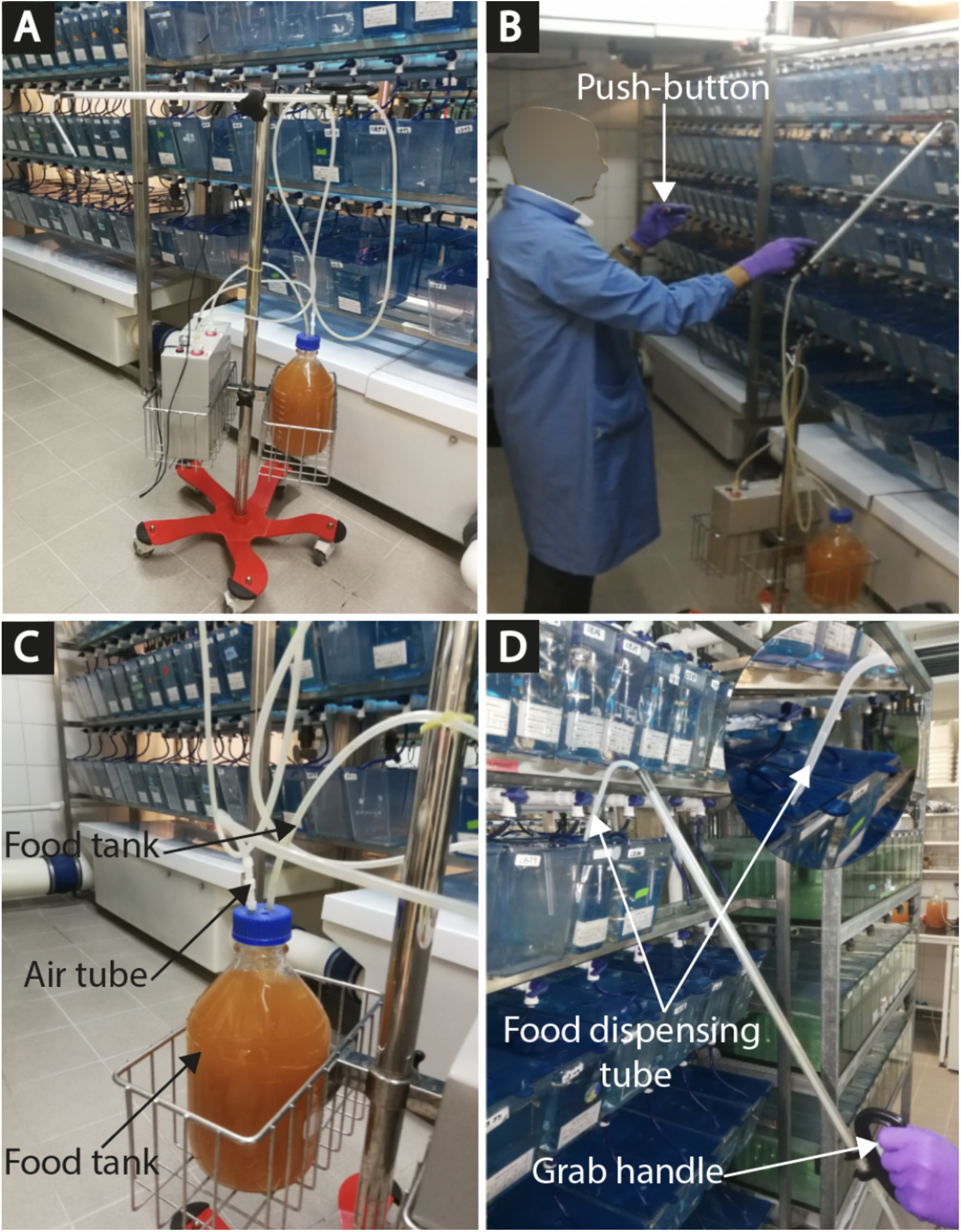
Live-food dispenser system. A) Full view of live-food dispenser system with wheeled basket cart - B) Staff from imaging facility feeds zebrafish - C) The food tank - D) Food dispenser tube.

**Figure 3:**
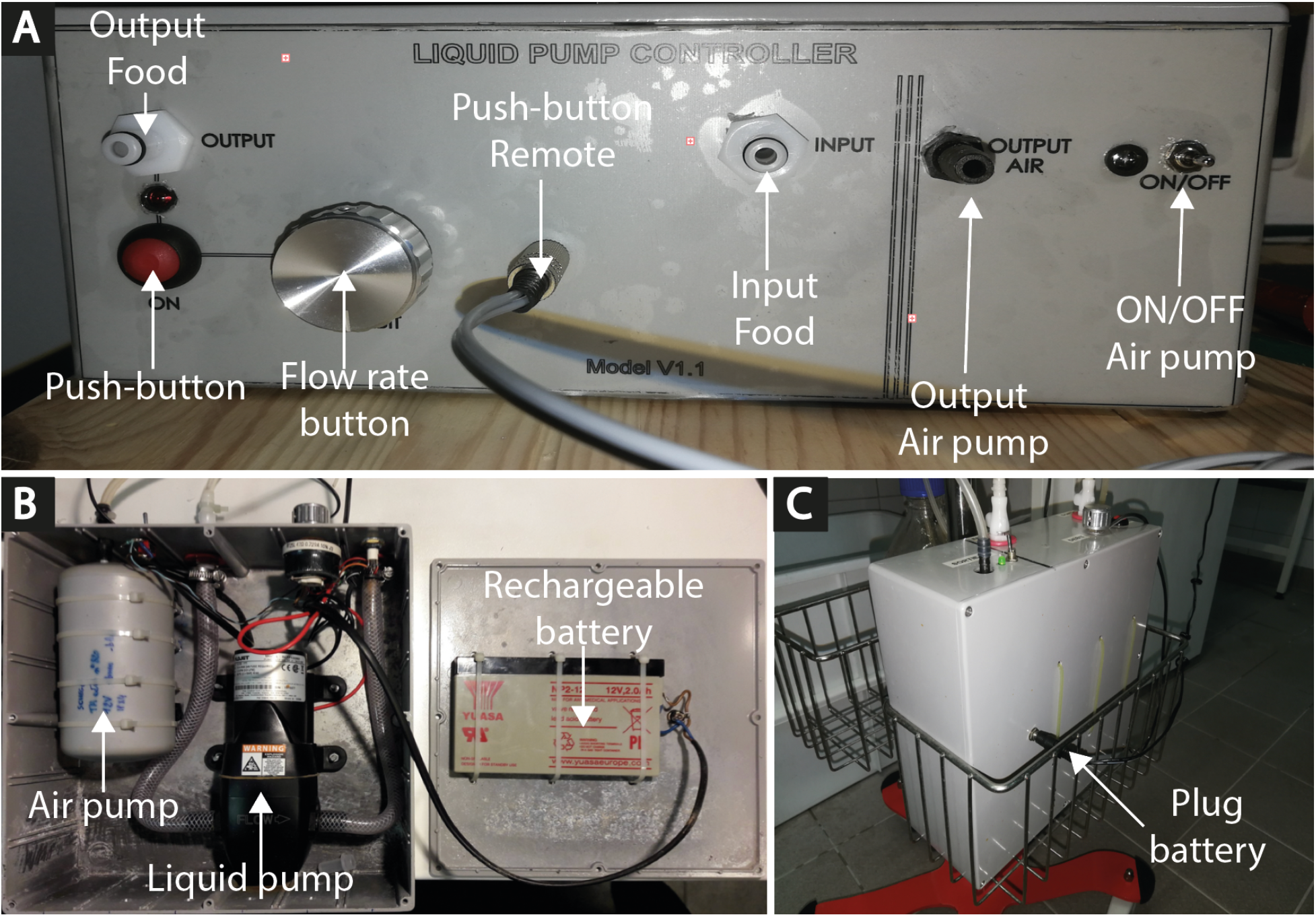
Electronics box. A) Front of view from electronic box - B) Inside vie of the electronics box - C) Picture of the electronics box showing the jack for the rechargeable battery.

The button is mounted on a 3D printed ergonomic holder. The pump will work as long as the button is pressed, thus the quantity of Artemia can be easily adjusted according to the number of fish in each aquarium (Figure 2B). The pump used is big enough (9.5mm horse diameter, diaphragm pump) to prevent any damage to the Artemia (other types of pumps tested damaged or killed the Artemia). After leaving the pump, the Artemia flow through a flexible tube connected to a rigid one (Figure 3).

The latter has a curved edge that enables feeding the upper aquariums of a rack without stepping on a footboard (Figure 2D). A second pump bubbles the solution containing the Artemia, preventing their precipitation and improving oxygenation (Figure 2C).

The system uses a 12 Volts rechargeable battery giving it freedom to move around the fish facility without the need of cable extensions, and more importantly is safer for the users (high voltage in a humid and wet environment can be dangerous for the users, Figure 3B). The Artemia reservoir, the pumps and the electronics box are all mounted on a very stable wheeled basket cart (see Figure 1A), allowing its simple and secure displacement around the fish facility.

All components can be found in Table 1. The wiring diagrams are described in Figure 4. A specific mounting protocol can be downloaded at www.fablab.biologie.ens.fr/live_food_system_download.html

**Table 1:**
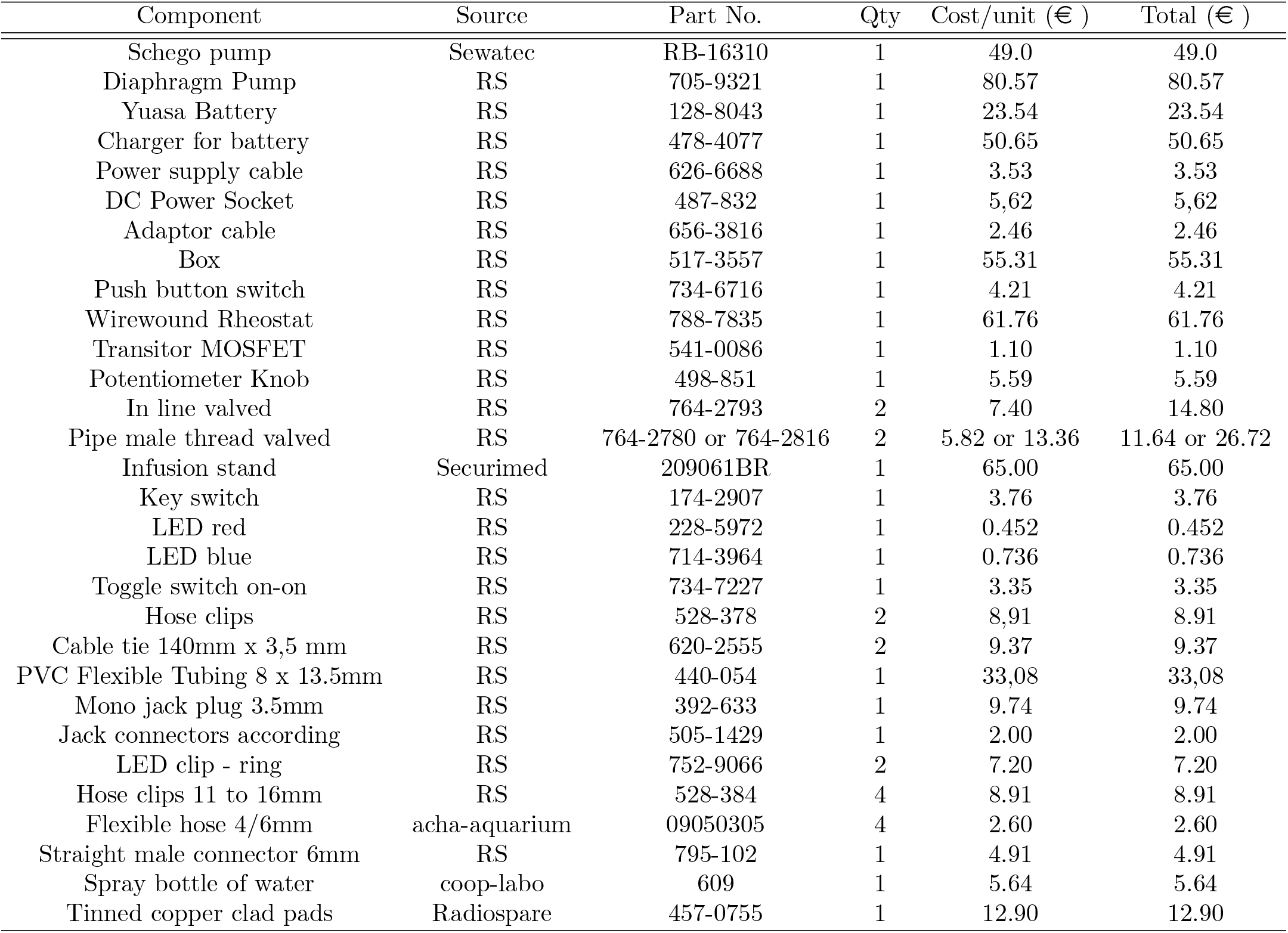
List of materials and components

| Component | Source | Part No. | Qty | Cost/unit (€ ) | Total (€ ) |
| --- | --- | --- | --- | --- | --- |
| Schego pump | Sewatec | RB-16310 | 1 | 49.0 | 49.0 |
| Diaphragm Pump | RS | 705-9321 | 1 | 80.57 | 80.57 |
| Yuasa Battery | RS | 128-8043 | 1 | 23.54 | 23.54 |
| Charger for battery | RS | 478-4077 | 1 | 50.65 | 50.65 |
| Power supply cable | RS | 626-6688 | 1 | 3.53 | 3.53 |
| DC Power Socket | RS | 487-832 | 1 | 5,62 | 5,62 |
| Adaptor cable | RS | 656-3816 | 1 | 2.46 | 2.46 |
| Box | RS | 517-3557 | 1 | 55.31 | 55.31 |
| Push button switch | RS | 734-6716 | 1 | 4.21 | 4.21 |
| Wirewound Rheostat | RS | 788-7835 | 1 | 61.76 | 61.76 |
| Transistor MOSFET | RS | 541-0086 | 1 | 1.10 | 1.10 |
| Potentiometer Knob | RS | 498-851 | 1 | 5.59 | 5.59 |
| In line valved | RS | 764-2793 | 2 | 7.40 | 14.80 |
| Pipe male thread valved | RS | 764-2780 or 764-2816 | 2 | 5.82 or 13.36 | 11.64 or 26.72 |
| Infusion stand | Securimed | 209061BR | 1 | 65.00 | 65.00 |
| Key switch | RS | 174-2907 | 1 | 3.76 | 3.76 |
| LED red | RS | 228-5972 | 1 | 0.452 | 0.452 |
| LED blue | RS | 714-3964 | 1 | 0.736 | 0.736 |
| Toggle switch on-on | RS | 734-7227 | 1 | 3.35 | 3.35 |
| Hose clips | RS | 528-378 | 2 | 8,91 | 8.91 |
| Cable tie 140mm x 3,5 mm | RS | 620-2555 | 2 | 9.37 | 9.37 |
| PVC Flexible Tubing 8 x 13.5mm | RS | 440-054 | 1 | 33,08 | 33,08 |
| Mono jack plug 3.5mm | RS | 392-633 | 1 | 9.74 | 9.74 |
| Jack connectors according | RS | 505-1429 | 1 | 2.00 | 2.00 |
| LED clip - ring | RS | 752-9066 | 2 | 7.20 | 7.20 |
| Hose clips 11 to 16mm | RS | 528-384 | 4 | 8.91 | 8.91 |
| Flexible hose 4/6mm | acha-aquarium | 09050305 | 4 | 2.60 | 2.60 |
| Straight male connector 6mm | RS | 795-102 | 1 | 4.91 | 4.91 |
| Spray bottle of water | coop-labo | 609 | 1 | 5.64 | 5.64 |
| Tinned copper clad pads | Radiospare | 457-0755 | 1 | 12.90 | 12.90 |

**Figure 4:**
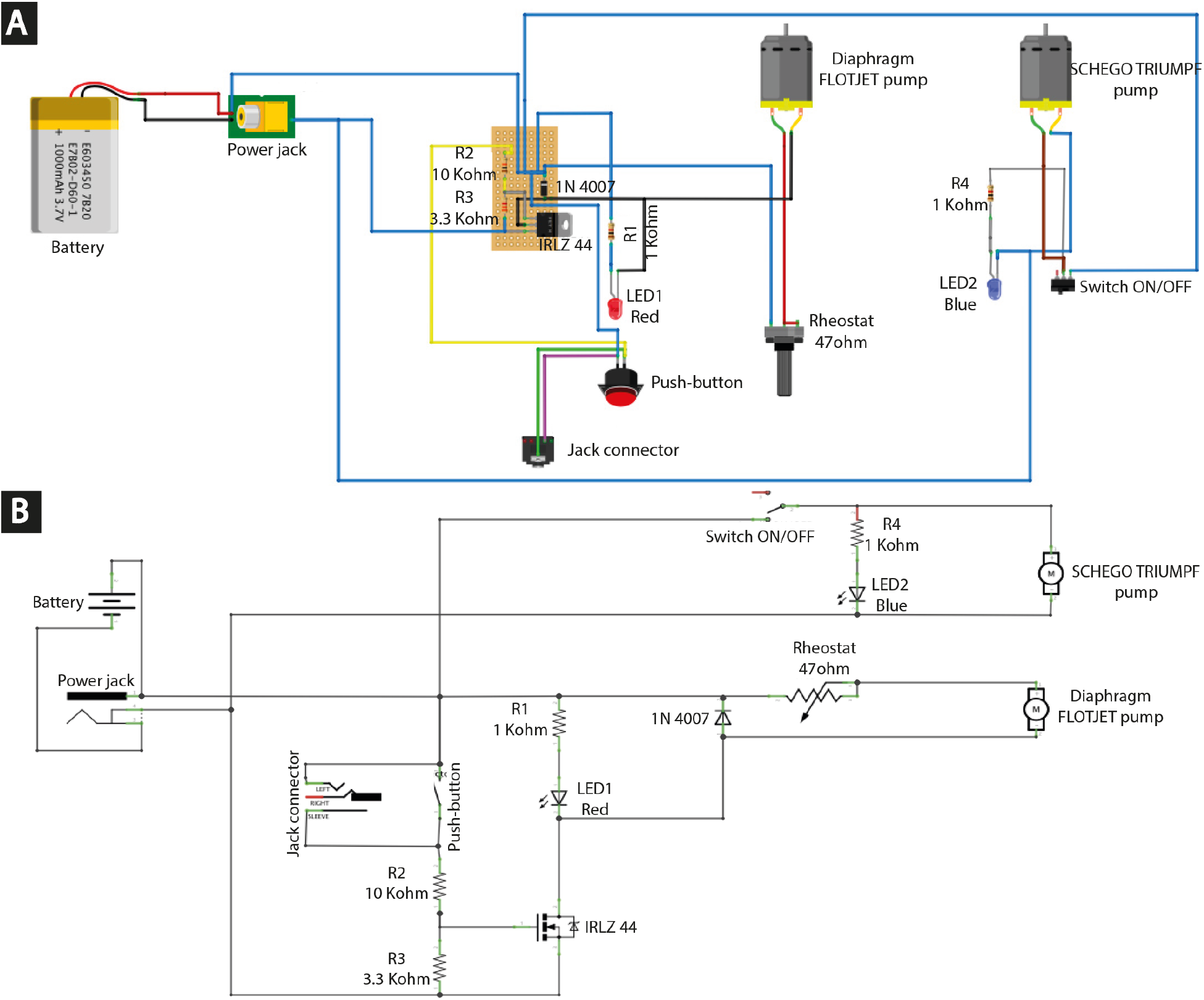
Electronics wiring. A) Diagram showing the pin configuration for the live-food dispenser system - B) Circuit diagram

In comparison to the time spent in feeding fish using a classic wash bottle, the live-food dispenser device significantly reduced the time spent in delivering Artemia to the 610 tanks of the fish facility (wash bottle: 36.9 min. live-food dispenser: 16.9 min., p=10^−10^, T-Test, n=7, Figure 11A). In addition, the variability (standard deviation, STD) in the weight of single doses was significantly smaller when using the live-food dispenser system (wash bottle STD: 1.5. live-food dispenser: 0.4. p=4^−14^, CHI Square Variance test, n=40, Figure 11C).

### Dry-food dispenser system

As a nutritional complement, zebrafish are also fed using dry food usually in the form of granules. Dry food is often delivered using forceps or pressure-based devices, requiring repetitive mechanical gestures.

To facilitate feeding and avoid these repetitive gestures, we designed a light-weight ergonomically conceived semi-automatic device capable of accurately delivering dry food.

This device is 3D printed (Figure 5) and uses standard electronic components (Table 2–4), except for the rotating and the fixed plate (Figure 6). These were built using a machining center (*Serrmac*) for better performance and durability.

**Figure 5:**
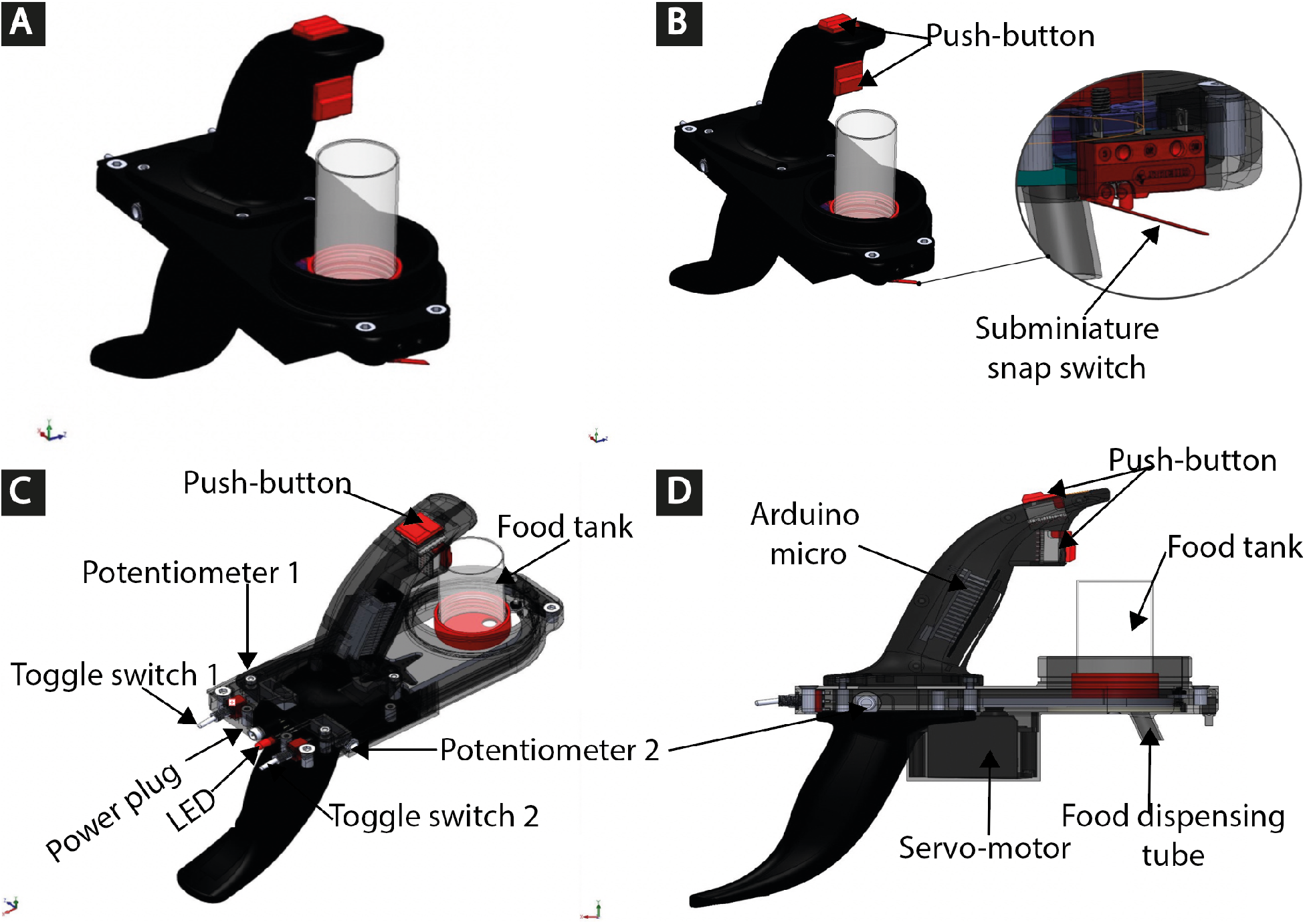
Dry-food dispenser system. A) 3D side view - B) 3D side view of buttons. Two buttons located on the joystick and a contact sensor that is activated as the device touches the lid of the aquarium. - C) 3D side view of switches. - D) Right side view.

**Table 2:** Material list parts

| Component | Source | Part No. | Qty | Cost/unit (€ ) | Total (€ ) |
| --- | --- | --- | --- | --- | --- |
| Polypropylene Bottles 60ml | Dutscher | 349035 | 3 | 0.30 | 0.90 |

**Table 3:** 3D print list parts

| Component | Source | Part No. | Qty | Cost/unit (€ ) | Total (€ ) |
| --- | --- | --- | --- | --- | --- |
| ABS filament | RS | 832-0359 | 1 | 30.99 | 30.99 |
| PVA filament | RS | 832-0494 | 1 | 49.63 | 49.63 |

**Table 4:** Electronic list parts

| Component | Source | Part No. | Qty | Cost/unit (€ ) | Total (€ ) |
| --- | --- | --- | --- | --- | --- |
| Servo-motor | RS | 781-3058 | 1 | 12.04 | 12.04 |
| Battery | RS | 614-2447 | 1 | 10.20 | 10.20 |
| Power cord | RS | 656-3816 | 1 | 2.46 | 2.46 |
| Charger for baterry | RS | 614-2447 | 1 | 54.17 | 54.17 |
| Power Socket | RS | 487-832 | 1 | 5,62 | 5,62 |
| Switch button | RS | 174-2907 | 2 | 3.76 | 7.52 |
| Arduino Micro | RS | 771-7667 | 1 | 20.79 | 20.79 |
| Toggle switch on-on | RS | 734-7227 | 2 | 3.22 | 6.44 |
| Tinned copper clad pads | RS | 457-0755 | 1 | 12.90 | 12.90 |
| Subminiature snap switch | RS | 868-1963 | 1 | 2.33 | 2.33 |
| Socket strip (5 way) | RS | 251-8200 | 4 | 0.63 | 2.52 |
| Socket strip (4 way) | RS | 251-81930 | 2 | 0.54 | 1.08 |
| Socket strip (3 way) | RS | 251-8187 | 2 | 0.47 | 0.94 |
| Socket strip (2 way) | RS | 251-8171 | 2 | 0.35 | 0.35 |
| PCB Header | RS | 547-3166 | 1 | 2.96 | 2.96 |
| Potentiometer 10KΩ | RS | 521-9041 | 2 | 1.53 | 3.06 |
| Potentiometer panel mount | RS | 107-6878 | 2 | 2.62 | 5.24 |
| Blue LED | RS | 235-9922 | 1 | 0.74 | 0.74 |
| Resistor 10KΩ | RS | 148-736 | 1 | 0.045 | 0.045 |
| Resistor 2.2KΩ | RS | 148-584 | 1 | 0.034 | 0.034 |
| Picth 3 | RS | 251-8092 | 1 | 1.13 | 1.14 |
| Screw M3 | RS | 478-743 | 4 | 26.98 | 26.98 |

**Figure 6:**
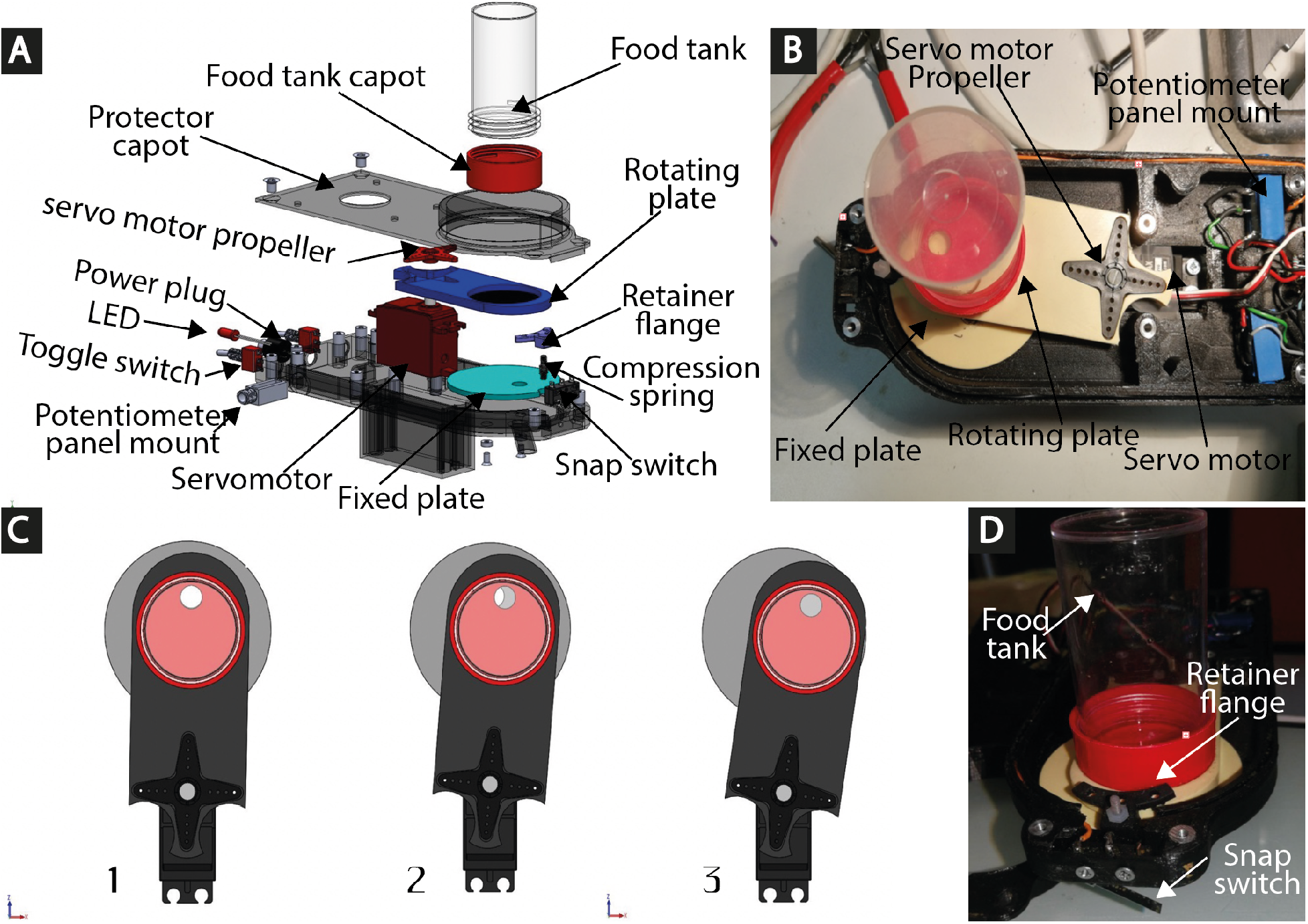
Dry-food feeder. A) Exploded view - B) Inside view showing the fixed and the rotating plates) - C) Examples of three possible moving positions of the dispenser (1) Position rest: no food is delivered - (2) Semi-open position: limited amount of food delivered - (3) Open Position: larger amount of food delivered relative to the semi-open position). - D) Front view.

However, a 3D printed version can also be used. It has two ergonomically designed handles to reduce torque, thus preventing arm fatigue and potential injuries (Figure 5).

Food delivery can be triggered using three different modes to avoid repetitive gestures: two sensitive buttons (only 1.5 Newton is required for activation) located at the natural proximity of the fingers and a contact sensor that is activated as the device touches the lid of the aquarium (Figure 5B and 7).

**Figure 7:**
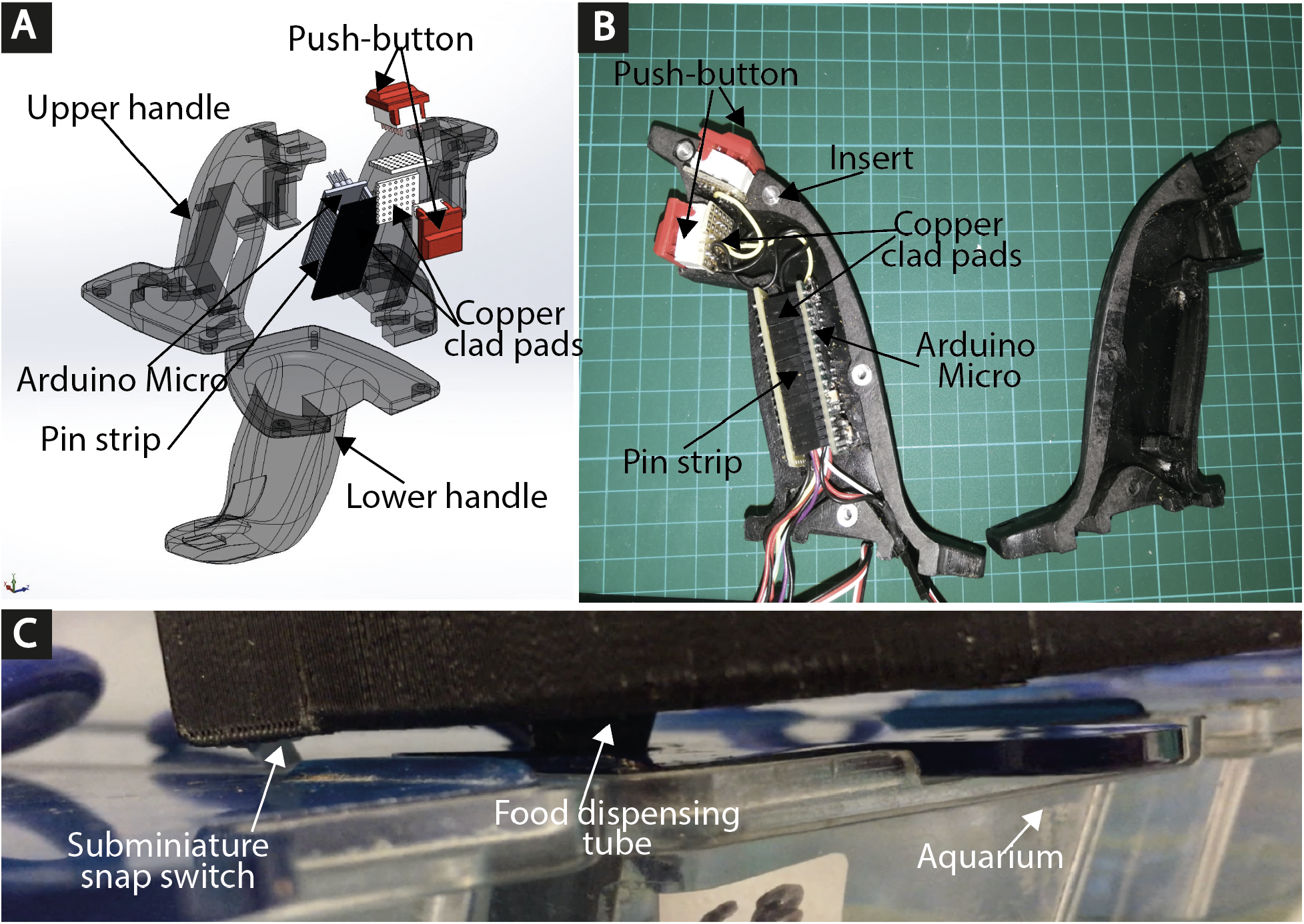
Push buttons. A) Exploded view of the upper handle - B) Inside view of the upper handle - C) Front button of the feeder that is activated as the device touches the lid of the aquarium.

The dry food is stocked in a standard Polypropylene 60 ml lab Bottles (*Dutscher 349035*, Figure 6A). Several of these standard reservoirs can be pre-filled and stocked in a refrigerator. These can then be easily screwed to the delivery system (Figure 6D). Upon activation of the device, the reservoir is displaced by a servomotor to momentary align it with 8 mm hole, allowing the liberation of the precise quantity of food (Figure 6C).

Two different types of food quantities can be pre-set. The quantity and the activation of the food delivery are controlled by an Arduino nano electronic board (Figure 8B).

**Figure 8:**
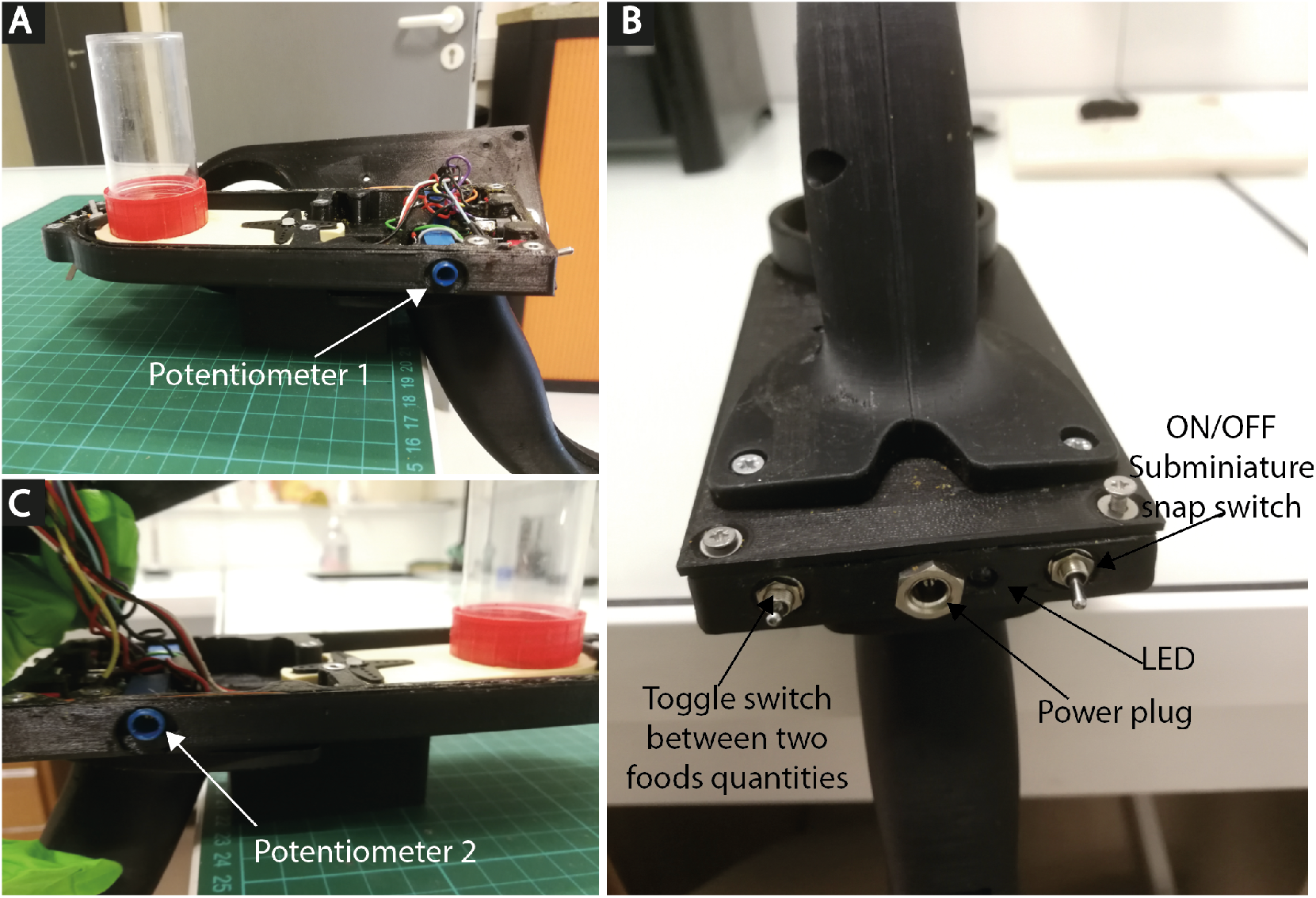
Electronic switches. A) Potentiometer 1 adjusts the quantity of food - B) Potentiometer 2 turns ON/OFF the snap switch - C) Back view with electronic toggle switch.

The whole system weights 950 grams, it measures 190×80×233 mm, and it is easy to build and to assemble (Figure 9). Figure 10 shows the Arduino code to control the device. A full assembly protocol can be downloaded at www.fablab.biologie.ens.fr/dry_food_system_download.html.

**Figure 9:**
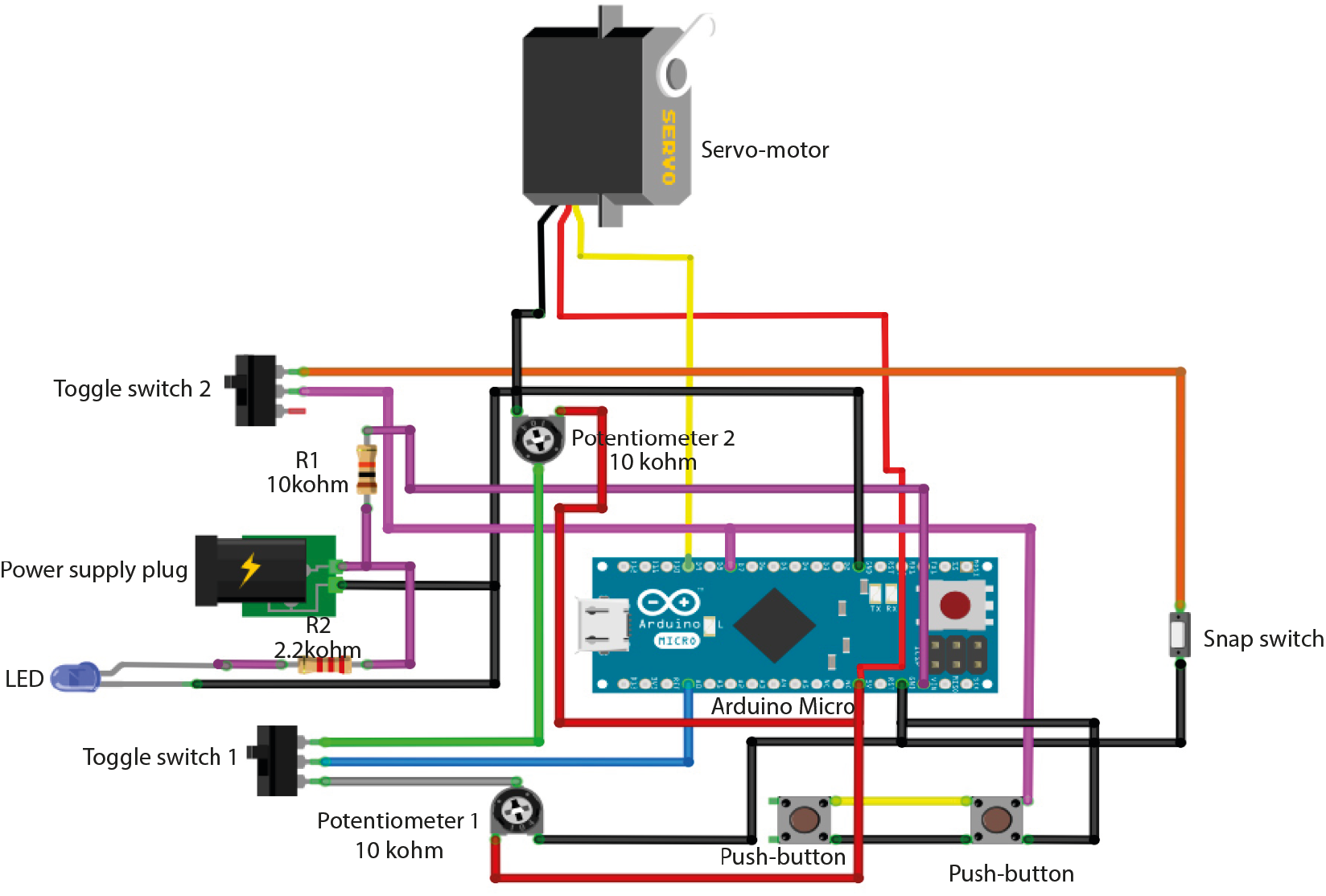
Visual circuit diagram

**Figure 10:**
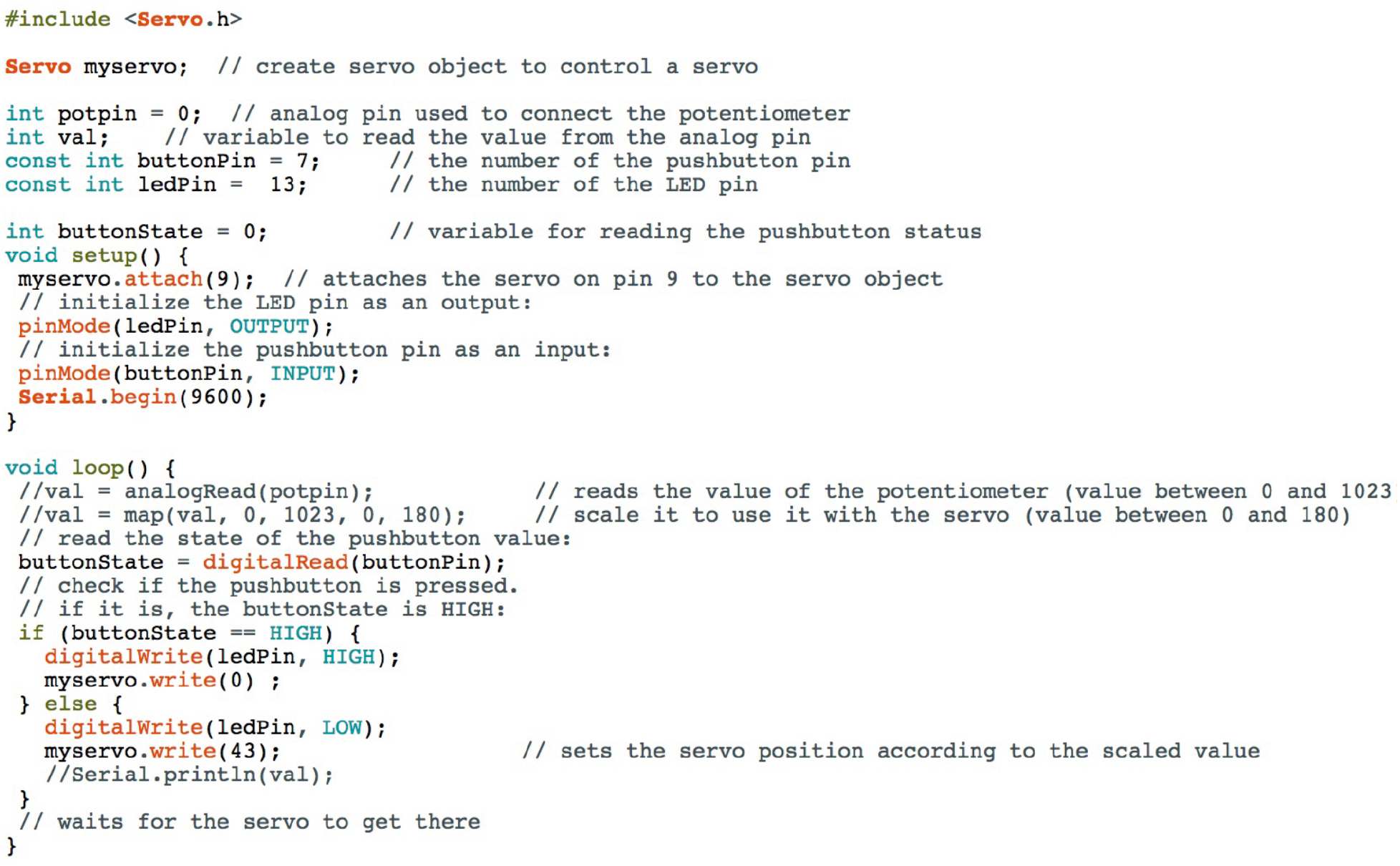
Arduino code

In comparison to the time spent in feeding fish using two fingers or a manual food dispenser (Special Diets Services, 824855), the dry-food dispenser device significantly reduced the time spent in delivering the dry food to the 610 tanks of the fish facility (hand: 39.4 min. manual dispenser: 33.8 min. dry-food dispenser: 16.3 min., p=7×10^−8^ and p=5×10^−7^, respectively, T-Test, n=7, Figure 11B). In addition, the variability (STD) in the delivered weight of single doses was significantly smaller when using the live-food dispenser system in comparison to the two-fingers method (hand: 0.06. dry-food dispenser: 0.009., p=4×10^−20^, CHI Square Variance test, n=40, Figure 11D). Although, the variance was smaller when comparing the dry-food dispenser with the manual food dispenser, this difference was not significant (manual dispenser: 0.01. dry-food dispenser: 0.09., p=0.16, CHI Square Variance test, n=40, Figure 11D).

**Figure 11:**
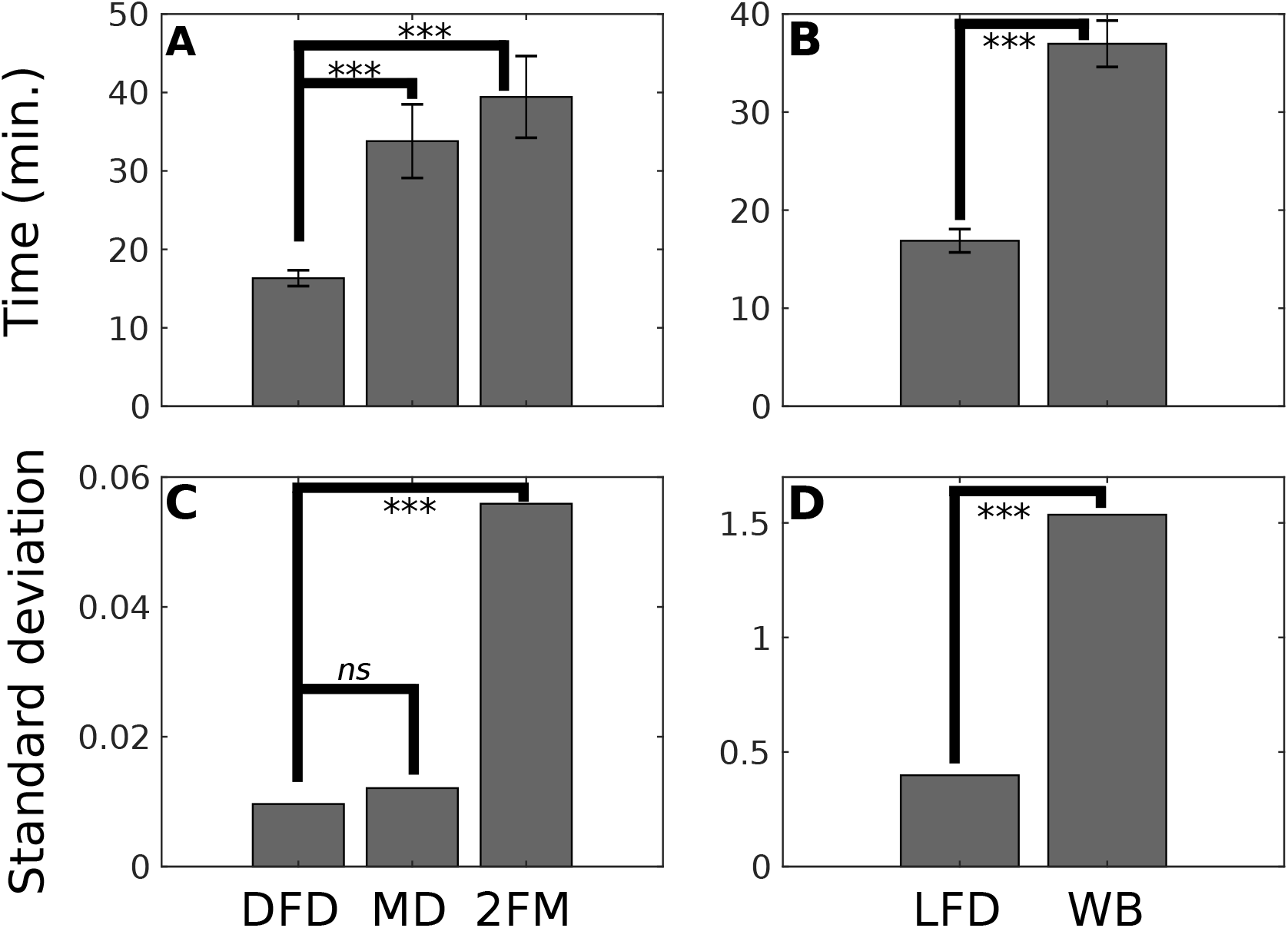
Comparison of feeding time and dose variance between the live- and dry-food delivery systems and conventional feeding methods. A) The time spent to feed fish in 610 tanks using Artemia. DFD: dry-food device, MD: manual device (Special food dispenser), 2FM: two fingers method. Error bars: standard deviation. B) As for (A), for the live-food device (LFD) and a common washing bottle (WB). Error bars: standard deviation. C) The standard deviation of the weight of single delivered doses (n=40). D) as for (C), for LFD and WB. ***: p <0.001. *ns*: non significant.

## Discussion

For the well being of zebrafish, they need to be fed several times a day with both dry and live food. This task demands repetitive motor gestures of the fish facility personnel which could, at long term, develop occupational disorders such as tendinitis or tenosynovitis [1, 2].

To prevent these occupational disorders and optimize fish feeding, we developed two ergonomic semiautomatic feeding systems for delivering dry and live food for small or mid-size fish facilities.

The use of these systems instead of the conventional manual methods fully healed the pathologies developed by our fish facility personnel and improved the speed and the accuracy of feeding.

These two systems are open source, economic and can be built by no experts using standard commercial components and 3D printing.

## Acknowledgments

We would like to thank doctor Véronique Sode and Marion Rahali (INSERM) for medical and ergonomic support. The authors also are grateful to Nathalie Cazenave and members of the Sumbre lab for comments and suggestions. The work was supported by the ERC CoG 726280 and the handicap pole of the INSERM.

